# *Pseudomonas aeruginosa* is capable of natural transformation in biofilms

**DOI:** 10.1101/859553

**Authors:** Laura M. Nolan, Lynne Turnbull, Marilyn Katrib, Sarah R. Osvath, Davide Losa, James J. Lazenby, Cynthia B. Whitchurch

**Affiliations:** The ithree institute, University of Technology Sydney, Ultimo, New South Wales, 2007, Australia; National Heart and Lung Institute, Imperial College London, London, SW3 6LR, UK; Microbes in the Food Chain Programme, Quadram Institute Bioscience, Norwich Research Park, Norwich, NR4 7UQ, UK; School of Biological Sciences, University of East Anglia, Norwich, NR4 7TJ, UK

## Abstract

Natural transformation is a mechanism that enables competent bacteria to acquire naked, exogenous DNA from the environment. It is a key process that facilitates the dissemination of antibiotic resistance and virulence determinants throughout bacterial populations. *Pseudomonas aeruginosa* is an opportunistic Gram-negative pathogen that produces large quantities of extracellular DNA (eDNA) that is required for biofilm formation. *P. aeruginosa* has a remarkable level of genome plasticity and diversity that suggests a high degree of horizontal gene transfer and recombination but is thought to be incapable of natural transformation. Here we show that *P. aeruginosa* possesses homologs of all proteins known to be involved in natural transformation in other bacterial species. We found that *P. aeruginosa* in biofilms is competent for natural transformation of both genomic and plasmid DNA. Furthermore, we demonstrate that type IV pili (T4P) facilitate but are not absolutely essential for natural transformation in *P. aeruginosa*.

## Introduction

The continued increase in antimicrobial resistance (AMR) levels is considered to be a significant global threat^1^. Horizontal gene transfer (HGT) is a key source of bacterial genome variation and evolution and is largely responsible for the acquisition of antibiotic resistance genes by bacterial pathogens^2^. Bacteria can acquire and heritably incorporate new genetic information via three HGT mechanisms: conjugation, transduction and natural transformation. Conjugation is a cell-contact dependent mechanism that transfers DNA directly from the cytoplasm of one bacterial cell into another. Transduction involves encapsidation of DNA into a bacteriophage which then injects the DNA into the recipient cell. The third HGT mechanism is natural transformation which involves the import of naked DNA from the environment through a specialised DNA transport apparatus^3,4^.

Extracellular DNA (eDNA) is present in significant quantities in both clinical and environmental settings, and provides a vast reservoir of genetic material that can be sampled by bacteria that are competent for natural transformation^5^. In many naturally competent bacterial species Type IV pili (T4P) are required for natural transformation^4^. While the exact role of T4P in natural transformation is unclear, the generally accepted model is that DNA binds to the pilus structure, which retracts and pulls the DNA to the cell surface. It is unclear whether or not DNA is translocated across the outer membrane through the PilQ secretin pore. The incoming DNA can then be accessed by the ComEA DNA translocation machinery in the periplasm, which mediates DNA uptake possibly by a ratchet mechanism^4,6^. In Gram-positive bacterial species that do not produce T4P, natural transformation involves a number of proteins with homology to T4P proteins which are thought to form a pseudopilus structure that spans the cell wall and is coupled to the DNA translocation complex at the cytoplasmic membrane^4,7^. Once exogenous DNA has been taken up by the cell it can be stably incorporated into the genome via recombination or transposition, or be maintained as an independent replicon if plasmid DNA is taken up by an appropriate host^7^. Natural plasmid transformation has also been described in *Escherichia coli*^8^. This appears to be a distinct mechanism which does not require homologs of the T4P/competence pseudopilus or DNA translocation machinery^9^.

*Pseudomonas aeruginosa* is a highly antibiotic resistant Gram-negative bacterium which is a part of the ‘ESKAPE’ group of pathogens that pose a serious health risk worldwide. *P. aeruginosa* readily acquires antibiotic resistance determinants, and demonstrates a high degree of genomic diversity and malleability similar to that seen in naturally transformable bacteria^10,11^. Despite this, *P. aeruginosa* has long been thought to be incapable of natural transformation^12^. *P. aeruginosa* is a model organism for studying T4P^13^. Interestingly, *P. aeruginosa* produces copious quantities of eDNA under conditions that promote T4P production such as in static broth cultures^14^, biofilms^15^ and during twitching motility-mediated biofilm expansion^16,17^. We therefore hypothesised that *P. aeruginosa* may be competent for natural transformation under conditions that promote both T4P expression and eDNA production. Here we show that some clinical and laboratory strains of *P. aeruginosa* are in fact capable of natural transformation under these conditions.

## Methods

### Strains, plasmids and growth conditions

*P. aeruginosa* strains used in this study were PAO1 (ATCC 15692), PAK^18^, PA14^19^, PA103^20^, PAO1_GFP_ which contains mini-Tn7-Gm^R^-P_A1/04/03_-*egfp* encoding *gfp* and *aac1* (Gm^R^) inserted downstream of *glmS*^21^, PAO1_CTX_ which contains miniCTX2 encoding *tet* (Tc^R^) inserted into the *attB* site of the chromosome^22^ and T4P mutants PAK*pilA*::*TcR*^23^, and Tn*5*-B21 mutants of *pilQ*^24^, *pilT*^25^, *pilV*^26^, *fimV*^27^. The *P. aeruginosa* CF sputum clinical isolates were obtained from David Armstrong at Monash Medical Centre (Melbourne, Australia), and the otitis externa *P. aeruginosa* clinical isolates were obtained from Di Olden at Gribbles Pathology Melbourne (Australia). The pUCPSK plasmid used is a non-conjugative *E. coli-P. aeruginosa* shuttle vector encoding *bla* which confers carbenicillin resistance (Carb^R^) in *P. aeruginosa*^28^. *E. coli* Dh5α (*recA, endA1, gyrA96, hsdR17, thi-1, supE44, relA1, φ80, dlacZΔM15*) was used as a host strain for pUCPSK and was miniprepped from *P. aeruginosa* and *E. coli* strains using a Qiagen miniprep kit according to manufacturer’s instructions.

*P. aeruginosa* was cultured on lysogeny broth (LB) solidified with agar at 1.5% (w/v) for routine maintenance and at 1.5% or 1% (w/v) for colony biofilm assays and grown in cation-adjusted Mueller Hinton Broth (CAMHB) at 37°C for all static broth and flow biofilm assays. Antibiotics were used at the following concentrations (w/v) as required: ampicillin 50 µg/ml for *E. coli* and carbenicillin 250 µg/ml, gentamicin 100 µg/ml and tetracycline 100 µg/ml for *P. aeruginosa*.

### Bioinformatics and data and statistical analyses

Homologs of proteins involved in natural transformation were identified in *P. aeruginosa* PAO1 using BLASTp^29^. The Pseudomonas.com resource^30^ and the PAK genome^31^ were used to identify *P. aeruginosa* orthologs.

Data was graphed and analysed using Graph Pad Prism version 8.0. The number of replicates and any statistical tests are described in Figure legends.

### Colony biofilm assay

Overnight cultures of *P. aeruginosa* were grown in 2 ml CAMHB at 37°C, shaking at 250 rpm. A 10 µL plastic loop was used to generate a 1 cm patch of the overnight culture on a dry 1% LBA plate. This was then incubated overnight at 37°C. The next day 10 µL of pUCPSK plasmid DNA (at a final concentration of 5 µg) was spotted onto the established colony biofilm and allowed to dry into the cells. The plate was then incubated with the agar downwards at 37°C for the indicated time. After incubation the colony biofilm was harvested from the plate into 1 mL LB, vortexed to resuspend and then incubated at 37°C for 30 min to fully resuspend the cells. The cell suspension was then spread plated between two 150 mm LBA plates with appropriate antibiotic selection and incubated for 24 hr at 37°C.

### Static broth assay

Overnight cultures of *P. aeruginosa* were grown in 2 ml CAMHB at 37°C, shaking at 250 rpm. 40µL of overnight culture was subcultured into 2 ml fresh CAMHB with DNA added at the indicated concentration. The media, cells and DNA were then mixed and incubated at 37°C statically for 24 hr. Note for the shaking broth assay the same setup was used however the culture was incubated with shaking at 250 rpm. In both cases after incubation the cell suspension was then spread plated between two 150 mm LBA plates with appropriate antibiotic selection and incubated for 24 hr at 37°C.

To image the aggregates present in static broth cultures *P. aeruginosa* was grown as described for the static broth assay in a glass-bottomed Ibidi® μ-Dish and visualised using DeltaVision Elite inverted research microscope with a × 100 1.4 numerical aperture UPlanFLN objective, InsightSSI illumination, SoftWorX acquisition software, fitted with a WeatherStation environmental chamber (Applied Precision, GE Healthcare, Issaquah, WA, USA) and a scientific CMOS 15-bit camera (pco.edge, PCO AG, Kelheim, Germany).

### Isolation of DNA for use in continuous flow biofilm assays

Chromosomal DNA (gDNA) was purified from PAO1Tn7::*gfp-aac1* cells using the Epicentre® Masterpure DNA purification kit. Extracellular DNA (eDNA) was purified from a confluent lawn of PAO1Tn7::*gfp-aac1* cultured overnight on MacConkey agar containing 5% (v/v) glycerol. Bacteria were suspended in sterile phosphate buffered saline (PBS), centrifuged and the supernatant filtered through 0.2 μm PES membrane. eDNA present in the supernatant was further purified by removal of proteins and ethanol precipitation as reported previously^32^. Sterility of all DNA samples was confirmed prior to use by plating onto LB agar.

### Continuous flow biofilm assays

10 cm lengths of Tygon laboratory tubing (2mm ID) were inoculated with 1/100 dilution of an overnight culture of *P. aeruginosa* in CAMHB and allowed to attach for 2 hours under static conditions after which continuous flow was commenced at a rate of 80 μL/min at room temperature. Influent media was CAMHB containing either no added DNA or DNA added at a final concentration of 1 μg/mL for pUCPSK plasmid DNA, or 0.5 mg/mL for gDNA or eDNA. At harvest, the attached biofilm was removed by sonication and biofilm-associated bacteria collected by centrifugation. Transformants were selected by plating onto LB agar with appropriate antibiotic selection and incubated for 24 hr at 37°C. To visualise natural transformation by PAO1 continuous flow biofilms, an IBIDI® μ-slide I (with flow kit) was inoculated and cultured as described for the Tygon tubing biofilms. Biofilms were imaged using an Olympus IX71 inverted research microscope with a × 100 1.4 numerical aperture UPlanFLN objective, FViewII monochromatic camera and AnalySIS Research acquisition software (Olympus Australia, Notting Hill, VIC, Australia) fitted with an environmental chamber (Solent Scientific, Segensworth, UK) for fluorescent imaging.

### Confirmation of natural transformation events from continuous flow biofilms

The presence of mini-Tn7-Gm^R^-P_A1/04/03_-*egfp* at the chromosomal attTn7 site was confirmed by PCR using primers Tn7_-up_ (5’CGTATTCTTCGTCGGCGTGAC3’) and Tn7_-down_ (5’CGAAGCCGCCGACAAGGG3’). Expression of GFP was confirmed by epifluorescence microscopy on an Olympus IX71. The presence of mini-CTX2 at the chromosomal *attB* site was confirmed by PCR using primers P_ser-up_ (5’CGAGTGGTTTAAGGCAACGGTCTTGA3’) and P_ser-down_ (5’AGTTCGGCCTGGTGGAACAACTCG 3’)^22^. To confirm the presence of pUCPSK in *P. aeruginosa*, plasmid DNA was extracted from *P. aeruginosa*, transformed into *E. coli*, extracted and confirmed by sequencing with M13_-FUP_ (5’TGTAAAACGACGGCCAGT3’).

## Results

Bioinformatic analyses of the sequenced *P. aeruginosa* strains PAO1, PA14 and PAK show that each of these strains encode homologs of all genes known to be involved in natural transformation in other bacterial species (Table 1). To determine if *P. aeruginosa* might be capable of natural transformation in biofilms, we established biofilms with a 1:1 mixture of PAO1_GFP_ (Gm^R^) and PAO1 [pUCPSK] (Carb^R^) under continuous flow conditions. Biofilm effluent was collected each day for 4 days and bacteria tested for their ability to grow on LB agar plates containing both gentamicin and carbenicillin. An average of 50-100 Gm^R^/Carb^R^ colonies (resistant to both gentamicin and carbenicillin) were obtained from the effluent of mixed PAO1_GFP_ and PAO1 [pUCPSK] biofilms on day 1 and confluent lawns of Gm^R^/Carb^R^ colonies obtained from day 2 onwards. The presence of mini-Tn7-Gm^R^-P_A1/04/03_-*egfp* at the chromosomal *attTn7* site in these colonies was confirmed by PCR amplification. To confirm that these colonies also possessed pUCPSK, plasmid DNA was extracted, transformed into *E. coli* and confirmed by sequencing. Neither the PAO1_GFP_ or PAO1 [pUCPSK] strains used to establish these mixed biofilms, or effluent from control single strain biofilms were able to grow on the dual antibiotic selection plates. As PAO1 lacks a prophage capable of transduction and pUCPSK is a non-conjugative plasmid, these results suggest that HGT of plasmid DNA and/or chromosomal DNA might occur via natural transformation in *P. aeruginosa* biofilms.

**Table 1.** Homologs of proteins involved in natural transformation in a range of bacteria.

| Competence Protein | *Bacillus subtilis* | *Streptococcus pneumoniae* | *Haemophilus influenzae* | *Thermus thermophilus* | *Pseudomonas stutzeri* | *Neisseria gonorrhoeae* | *Pseudomonas aeruginosa* * |
| --- | --- | --- | --- | --- | --- | --- | --- |
| **T4P/Competence Pseudopilus** |  |  |  |  |  |  |  |
| Traffic NTPase(s) | ComGA | ComGA | PilB | PilF | PilT, PilU | PilF, PilT | PilB, PilT, PilU |
| Polytopic membrane protein | ComGB | ComGB | PilC | PilC | PilC | PilG | PilC |
| Pilins or pseudopilins | ComGC, -GD, -GE, -GG | CglC, CglD | PilA | PilA1, -A2, -A3, -A4 | PilA1 | PilE, ComP | PilA, -V, -W, -X, -E, FimT, FimU |
| Prepilin peptidase | ComC | CilC | PilD | PilD |  | PilD | PilD |
| Secretin/pilot | na | na | ComE | PilQ |  | PilQ/PilP | PilQ/PilP |
| **DNA translocation machinery** |  |  |  |  |  |  |  |
| DNA receptor | ComEA | ComEA |  | ComEA |  | ComE | PA3140 |
| Membrane channel | ComEC | ComEC | Rec-2 | ComEC | ComA | ComA | PA2984 |
| ATP-binding protein | ComFA | ComFA |  |  | ExbB |  | PA2983 |
| **Other** |  |  | DprA (Smf) |  |  |  | PA0021 |
|  |  |  | TfoX (Sxy) |  |  |  | PA4703 |
|  |  |  | CRP |  |  |  | Vfr |
|  |  |  | CyaA |  |  |  | CyaA, CyaB |
|  |  |  | ComM |  |  |  | PA5290 |
|  |  |  | ComF |  |  |  | PA0489 |

\*Using *P. aeruginosa* PAO1 gene nomenclature

To determine if HGT of chromosomal DNA did indeed occur via natural transformation, and the above result was not just due to HGT of plasmid DNA, we established biofilms with a 1:1 mixture of PAO1_GFP_ (Gm^R^) and PAO1_CTX_ (Tc^R^) under continuous flow conditions. Biofilm effluent was collected each day for 8 days and bacteria tested for their ability to grow on LB agar plates containing both gentamicin and tetracycline. Whilst no Gm^R^/Tc^R^colonies were obtained from days 1-4, from days 5-8 an average of 3 Gm^R^/Tc^R^ colonies that were resistant to both antibiotics and expressed GFP were recovered per day. The presence of both mini-Tn7-Gm^R^-P_A1/04/03_-*egfp* and mini-CTX2 in these colonies was confirmed by PCR. Importantly, neither the PAO1_GFP_ or PAO1_CTX_ strains used to inoculate the mixed biofilms or effluent from control single-strain biofilms were able to grow on the dual antibiotic selection plates. As neither conjugation or transduction is likely to account for these HGT events in PAO1 biofilms, these results suggest that *P. aeruginosa* is able to acquire and incorporate chromosomal DNA and plasmids encoding antibiotic resistance genes via natural transformation in biofilms.

To confirm that *P. aeruginosa* is indeed capable of natural transformation and to rule out any possibility of HGT through transduction or conjugation, we performed a series of experiments to follow the uptake of purified, sterile exogenous DNA. We firstly examined plasmid DNA uptake under continuous flow biofilm conditions. PAO1 flow biofilms were cultured in the presence or absence of purified pUCPSK plasmid DNA (at a final concentration of 1 μg/mL) and the amount of natural transformation within the biofilm biomass and effluent assessed at days 3, 4 and 5. Carb^R^ colonies were obtained from both the biofilm biomass and biofilm effluent (Fig. 4a). No Carb^R^ colonies were obtained in the no DNA control.

**Figure 1.**
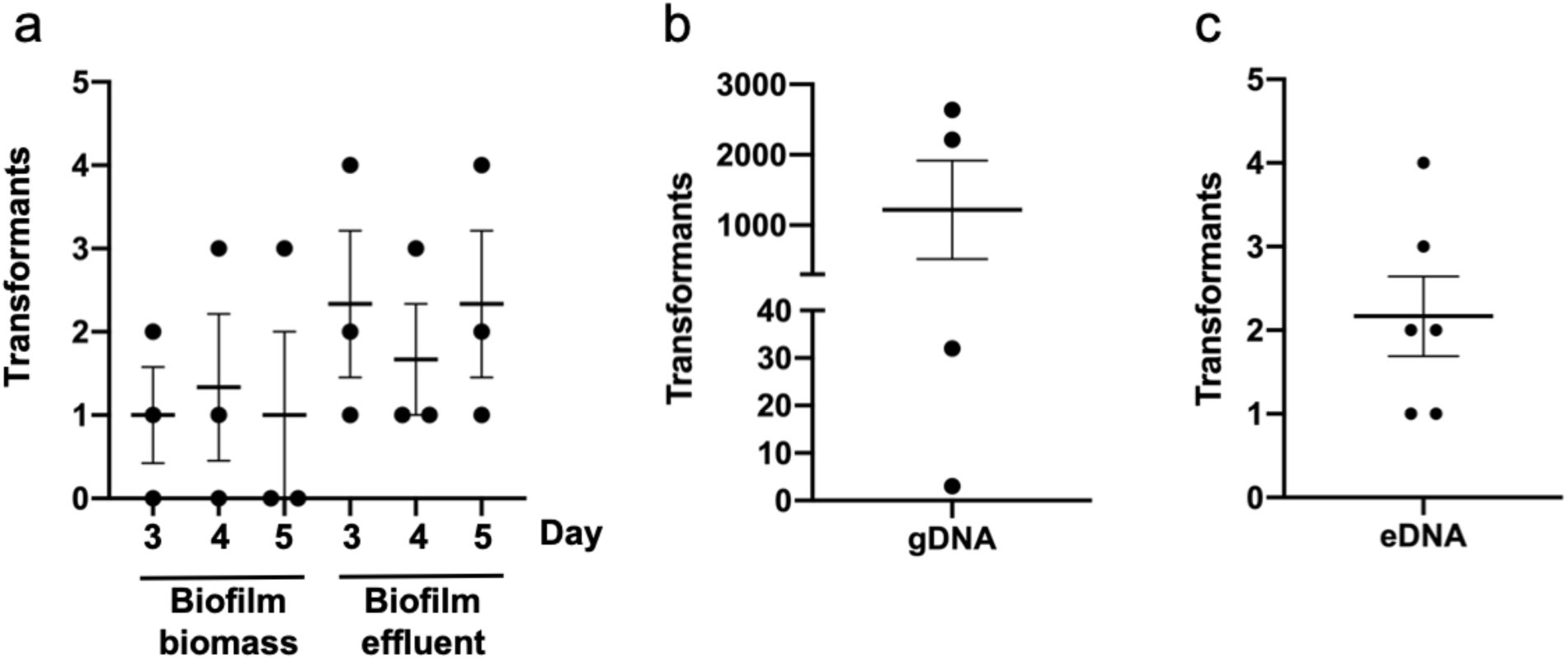
*P. aeruginosa* is capable of natural transformation in continuous flow biofilms. Carbenicillin or gentamicin resistant transformants obtained from continuous flow biofilms of PAO1 in Tygon tubing. For (a) pUCPSK was added to the media influent (at a final concentration of 1 μg/mL) with either the biofilm biomass or effluent harvested and plated on carbenicillin selective agar plates on the indicated day. For (b-c) sterile gDNA or eDNA obtained from PAO1_GFP_ (Gm^R^) (at a final concentration of 0.5 mg/mL) was added to the media influent and the biofilm biomass harvested and plated on gentamicin selective agar plates after 5 days. The mean of each set of technical triplicates was calculated to give an n≥3 which is presented as mean ± S.E.M.

**Figure 2.**
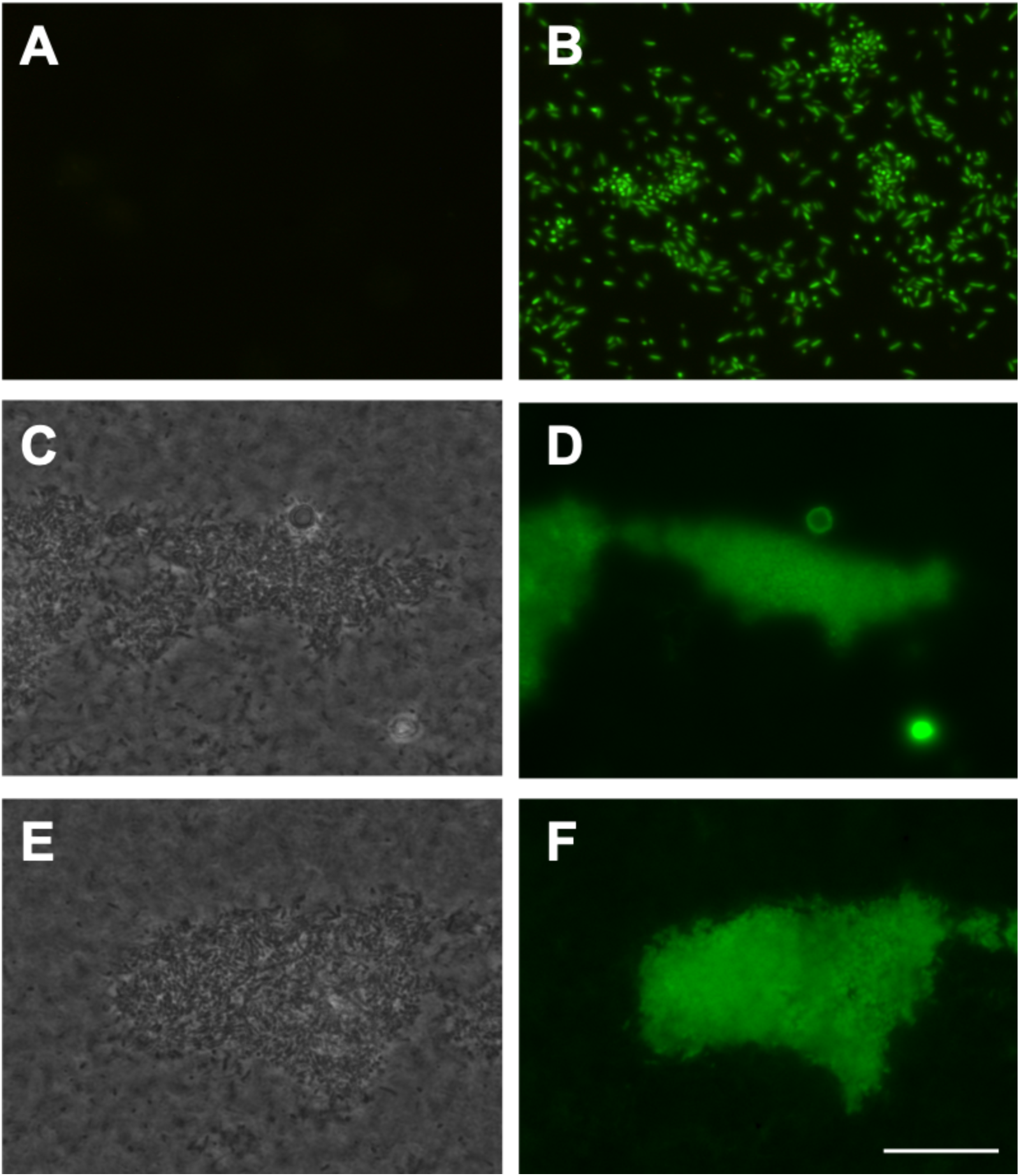
*P. aeruginosa* is capable of natural transformation of exogenous chromosomal DNA. The inoculum PAO1 strain did not express GFP (a). Sterile PAO1_GFP_ gDNA (at a final concentration of 0.5 mg/mL) was added to the media influent of continuous flow biofilms of PAO1 in Tygon tubing (b) or flow cells (c-f) and incubated for 5 days. Gentamicin resistant (Gm^R^) colonies obtained from Tygon tubing biofilms were resuspended in PBS and visualized by epifluorescence microscopy which showed all cells from Gm^R^ colonies expressed GFP (B). The biofilm biomass from PAO1 flow cells was visualized by phase contrast (c, e) or epifluorescence microscopy (d, f) which showed the presence of representative biofilm microcolonies expressing GFP. Scale bar 100 μm.

**Figure 3.**
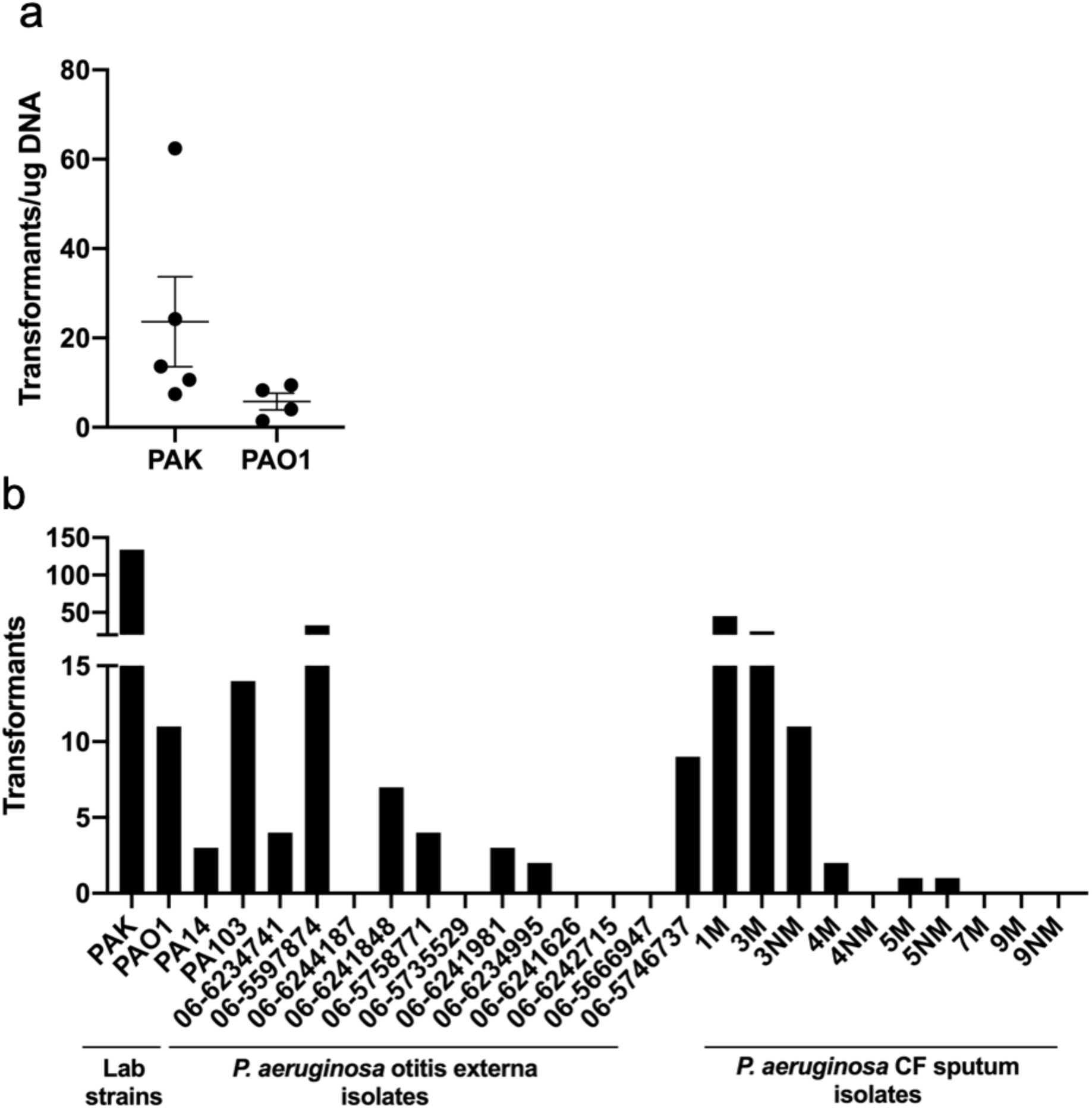
Lab and clinical strains of *P. aeruginosa* are capable of natural transformation of plasmid DNA within colony biofilms. pUCPSK (5 μg) DNA was applied to colony biofilms of (a) *P. aeruginosa* PAK or PAO1, or (b) *P. aeruginosa* lab or clinical strains and incubated for 2 hr. Cells were then harvested and the number of carbenicillin resistant transformants determined by spread plating on selective media. For (a) the mean of each set of technical triplicates was calculated to give an n≥4 which is presented as mean ± S.E.M. For (b) the values presented are from n=1. For the CF sputum isolates the designation M refers to mucoid phenotype, NM is non-mucoid.

**Figure 4.**
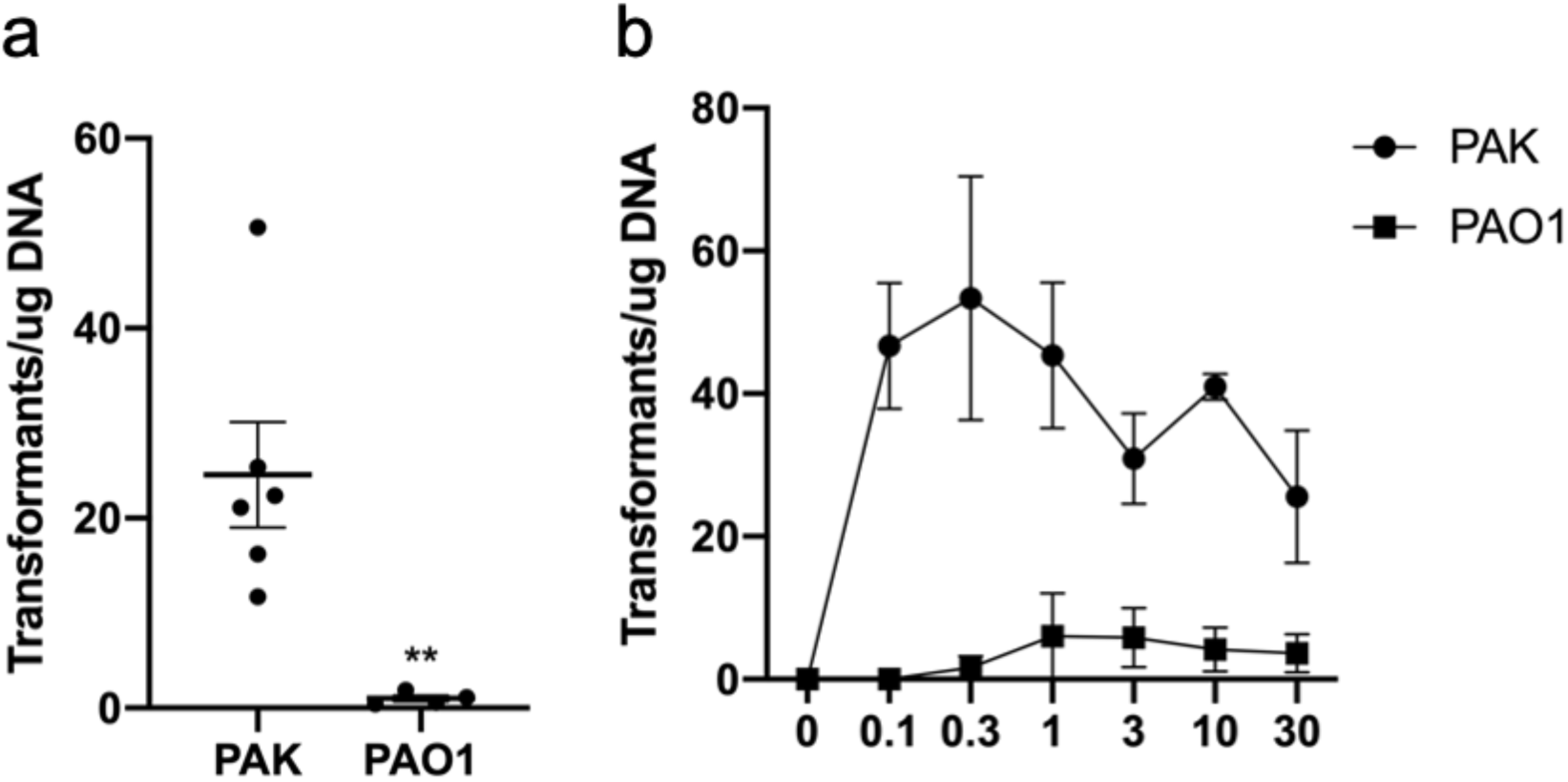
*P. aeruginosa* is capable of natural transformation of plasmid DNA in static broth cultures. Carbenicillin resistant transformants obtained from static broth cultures incubated for 24 hr with (a) 10 μg pUCPSK or (b) 0-30 μg pUCPSK. The mean of each set of technical triplicates was calculated to give an n≥3 which is presented as mean ± S.E.M. For (a) ** *P* <.005; Mann-Whitney *U*-test compared to PAK.

We also examined if natural transformation by uptake of exogenous chromosomal DNA occurs in biofilms cultured under continuous flow. *P. aeruginosa* PAO1 flow biofilms were cultured in the presence and absence of sterile gDNA or eDNA obtained from PAO1_GFP_ (Gm^R^) (at a final concentration of 0.5 mg/mL) in the media influent. After 5 days the number of Gm^R^ colonies recovered from the biofilm biomass were counted. This revealed extremely variable rates of natural transformation of gDNA by cells within the biofilm biomass across multiple experiments (Fig. 1b). This is not unexpected as the rate is likely to be dependent upon the time at which the natural transformation event occurred. If this event occurred early in the assay, we would expect many transformants recovered due to proliferation of the transformed cells. However, if transformation occurred later we would expect far fewer transformants as these did not have as long to proliferate. For the eDNA experiments, while some Gm^R^ transformants were obtained (Fig. 1c), the rate of natural transformation was overall much lower than for gDNA (Fig. 1b). This may be due to the integrity of the DNA as we observed via agarose gel electrophoresis that the eDNA used in these experiments was quite degraded compared with the gDNA, presumably through the action of nucleases present in the extracellular milieu. No Gm^R^ colonies were obtained for continuous flow biofilms in the absence of gDNA or eDNA indicating that the gentamicin resistant cells recovered from these assays was due to the presence of the exogenous chromosomal DNA. To further rule out the possibility of spontaneous resistance, the presence of the mini-Tn7-Gm^R^-P_A1/04/03_-*egfp* at the chromosomal attTn7 site in the biofilm-derived Gm^R^ colonies was confirmed by PCR. The presence of the *gfp* gene in the Gm^R^ colonies was also confirmed by visualisation of GFP expression using epifluorescence microscopy (Fig. 2b). No GFP expression was observed in the PAO1 inoculum strain (Fig. 2a). As it was not possible to directly visualise biofilms cultured in Tygon tubing, PAO1 continuous flow biofilms were cultured in transparent flow cells over 5 days in the presence and absence of gDNA obtained from PAO1_GFP_. Epifluorescence microscopy revealed microcolonies of GFP-expressing bacteria within the biofilm (Fig. 2c-f). No GFP expression was observed in the no DNA control biofilms.

To further investigate natural transformation by *P. aeruginosa* we set out to determine if this process with also occurring in colony biofilms on agar plates. We first grew wildtype strains PAK and PAO1 on LB agar overnight to form a colony biofilm on the surface of the agar. We then added 5 μg of sterile DNA (or the equivalent volume of sterile water) of the plasmid pUCPSK, onto the surface of the colony biofilm. After 2 hr incubation at 37°C the colony was resuspended and cells plated onto media containing carbenicillin to select for transformants that had acquired the plasmid. Only colony patches that had been exposed to plasmid DNA yielded Carb^R^ colonies (Fig. 3a), whereas colony patches exposed to sterile water yielded none. The transformation efficiency for PAK and PAO1 was 24.7 ± 10.1 and 5.8 ± 1.9 transformants/μg plasmid DNA, respectively. To confirm that the carbenicillin resistant colonies had acquired pUCPSK, plasmid DNA was extracted, re-transformed into *E. coli* and confirmed by sequencing. These observations indicate that a proportion of cells within colony biofilms of *P. aeruginosa* are competent for natural transformation and are able to take up and maintain plasmid DNA.

Given that both *P. aeruginosa* strains PAK and PAO1 appeared to be naturally competent, we wanted to determine if this was also the case for two other commonly utilised lab strains (PA14 and PA103) and clinical isolates. All *P. aeruginosa* strains were first confirmed to be carbenicillin sensitive prior to use. Thirteen *P. aeruginosa* otitis externa and twelve cystic fibrosis (CF) lung sputum isolates were assayed for the ability to uptake pUCPSK plasmid DNA in a colony biofilm. Of these, 7/12 otitis externa and 6/10 CF isolates were able to uptake exogenous plasmid DNA (Fig. 3b). No Carb^R^ colonies were obtained in the no plasmid DNA controls for each strain. Interestingly, transformation ability varied amongst both clinical and lab strains. Of the lab strains, PA14 was the least capable of natural transformation with PAK the most efficient. These results demonstrate that many lab and clinical isolate strains of *P. aeruginosa* are capable of natural transformation within colony biofilms, albeit with different efficiencies.

*P. aeruginosa* also expresses T4P when cultured in static nutrient broth^13^. Under these conditions *P. aeruginosa* forms biofilms and suspended microcolony aggregates that contain eDNA^14,33^. To determine if natural transformation also occurred in static broth cultures, 10 μg of sterile pUCPSK plasmid DNA (or the equivalent volume of sterile water) was added to a subculture of *P. aeruginosa* wildtype PAK or PAO1 and incubated statically at 37°C for 24 hrs. Cells were then recovered and plated onto media containing carbenicillin to select for transformants. Carb^R^ colonies were obtained for both PAO1 and PAK under these conditions, whereas no Carb^R^ colonies were identified in the water control. As was observed with colony biofilm transformations (Fig. 3a), PAK was more efficient for natural transformation of pUCPSK than PAO1 in static broth cultures, with efficiencies of 24.6 ± 5.5 and 1.0 ± 0.3 transformants/μg plasmid DNA, respectively (Fig. 4a).

To determine if transformation efficiency was dependent on the amount of DNA added, we performed static broth transformation assays with increasing amounts of plasmid DNA. Whilst we observed an increase in the number of transformants with increasing amounts of sterile plasmid DNA added (Fig. S1), the transformation efficiency (transformants/μg DNA) was relatively unchanged over the range of DNA quantities used for transformation (0-30 μg) (Fig. 4b).

We were also interested in determining whether *P. aeruginosa* was also able to uptake exogenous chromosomal DNA and integrate this into the chromosome in static broth cultures. To examine this, chromosomal DNA from PAO1_GFP_ (Gm^R^) was purified from either a sterile whole cell lysate (gDNA) or the total sterile eDNA from confluent agar plate culture and 15 μg added to static broth cultures of PAK or PAO1 for 24 hr. Cells were recovered and cultured on agar containing gentamicin to select for transformants. These assays revealed that natural transformation of gDNA occurred at a low frequency for both PAK and PAO1 in static broth cultures (Fig. S2). No natural transformation with sterile eDNA was observed in static broth cultures for either PAK or PAO1 (Fig. S2). As mentioned above for eDNA in continuous flow biofilms, this may be due to the degraded nature of the eDNA compared to the gDNA as visualised by agarose gel electrophoresis. No gentamicin resistant colonies were obtained in the no DNA controls.

*P. aeruginosa* produces more T4P when cultured in static broth cultures than under shaking conditions^13^. We investigated the effects on natural transformation efficiency of culturing under static or shaking conditions and found that although some natural transformation was still observed under shaking conditions, more transformants were obtained with static culture conditions (Fig. 5a), consistent with a role of T4P in natural transformation. To directly examine the role of T4P in natural transformation of *P. aeruginosa*, we added 10 μg sterile pUCPSK plasmid DNA to static broths of PAK mutants defective in the production of the pilin subunit (*pilA*), in T4P assembly (*pilV, pilQ, fimV*) and in T4P retraction (*pilT*). Interestingly, all T4P mutants were capable of some natural transformation of pUCPSK, however a significant reduction in transformation efficiency compared to wildtype PAK was observed (Fig. 5b). No Carb^R^ colonies were identified in the no DNA controls. There was no apparent difference in the transformation efficiency of mutants which either didn’t have any surface-assembled T4P (*pilA, pilV, pilQ, fimV*) or were unable to retract T4P (*pilT*) (Fig. 5b).

**Figure 5.**
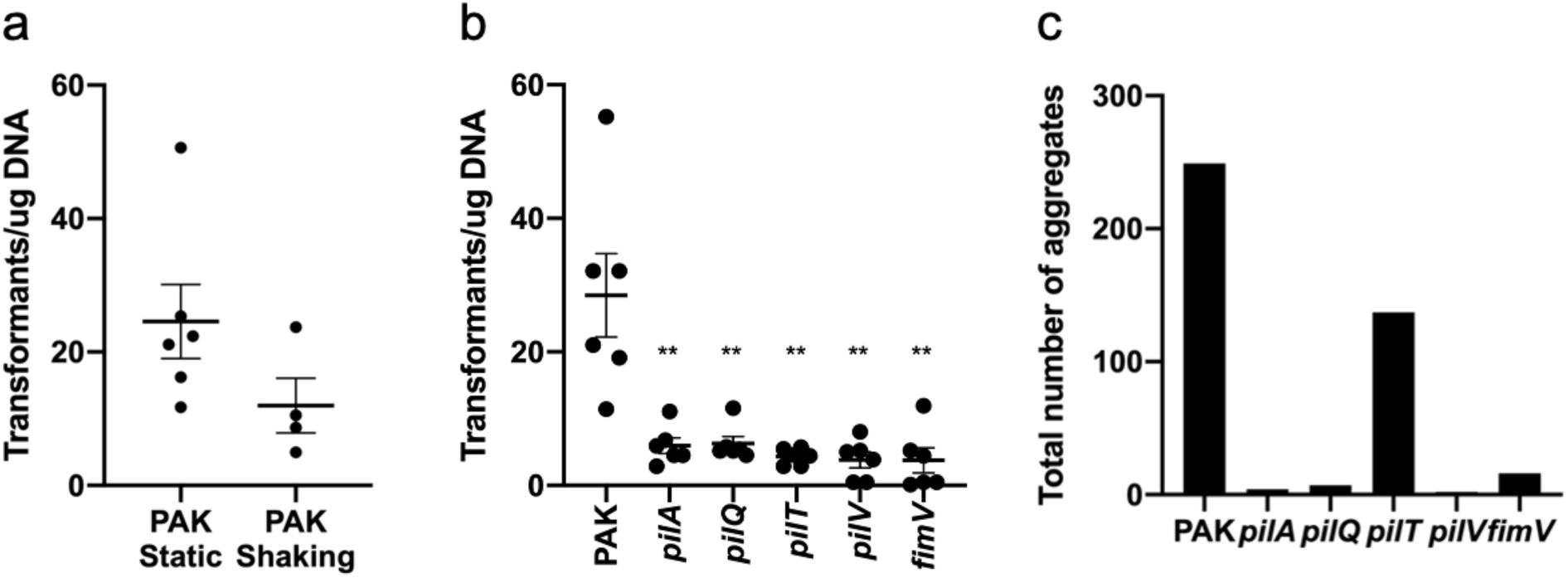
T4P are not absolutely required for natural transformation of plasmid DNA. Static (a-c) or shaking (a) broth cultures were incubated for 24 hr with 10 μg pUCPSK and the number of (a-b) carbenicillin resistant transformants or (c) aggregates observed determined. For (a-b) the mean of each set of technical triplicates was calculated to give an n≥3 which is presented as mean ± S.E.M. For (a) *P* > .05; Mann-Whitney *U*-test. For (b) ** *P* <.005; Mann-Whitney *U*-test compared to PAK. For (c) the total number of aggregates from 60 random fields imaged on 3 separate days (20 fields/day) is presented.

Given that wildtype *P. aeruginosa* forms aggregates in a static broth^14,33^ we considered the possibility that natural transformation was occurring within these aggregates in our static broth transformation assay. To determine if the reduction in transformation efficiency by T4P mutants (Fig. 5b) could be accounted for by the inability to form aggregates, we used microscopy to image wildtype PAK and T4P mutants under static broth conditions. This revealed that mutants unable to make surface-assembled T4P (*pilA, pilV, pilQ, fimV*) were also unable to form aggregates (Fig. 5c). A *pilT* mutant, which has a T4P retraction defect, was however still able to form aggregates (Fig. 5c). While the total number of aggregates was less than that observed for wildtype (Fig. 5c), this decrease does not account for the observed decrease in natural transformation efficiency (Fig. 5b). This suggests that it is the lack of functional T4P effects natural transformation and not other factors associated with aggregate formation.

Overall these data suggest that in *P. aeruginosa*, T4P facilitate transport of DNA to the cell surface but are not essential for natural transformation, at least of plasmid DNA. Furthermore, these observations indicate that during natural transformation in *P. aeruginosa* the PilQ secretin pore is not absolutely required for translocation of plasmid DNA across the outer membrane.

## Discussion

Here we have demonstrated, in contrast to current dogma, that *P. aeruginosa* is capable of natural transformation of both plasmid and chromosomal DNA under conditions that promote the expression of T4P and eDNA production, such as in static broth cultures and biofilms. We found that whilst T4P appear to be involved in facilitating plasmid DNA uptake, T4P are not absolutely required for natural transformation of plasmid DNA in *P. aeruginosa*. Furthermore, our data suggests that the PilQ secretin pore is not absolutely required for translocation of plasmid DNA across the outer membrane in this organism.

It is thought that separate uptake mechanisms for plasmid or chromosomal DNA are present in *E. coli*^8^. While natural transformation of chromosomal DNA and integration into the genome has not been demonstrated in *E. coli*, T4P/competence pseudopilus and DNA translocation machinery homologs have been shown to be responsible for uptake of chromosomal DNA for use as a nutrient source^34^. Natural plasmid transformation in *E. coli* does not utilise the competence pseudopilus uptake system although the precise mechanism for this process has not been determined^9^. At this stage we cannot rule out the possibility that there are two mechanisms in *P. aeruginosa* involved in the natural transformation of plasmid DNA or chromosomal DNA.

The finding that *P. aeruginosa* is capable of natural transformation is a paradigm shift in our understanding of how this pathogen acquires genetic diversity. Indeed, recombination has been identified as a major means of genetic diversity in *P. aeruginosa* cystic fibrosis (CF) lung isolates although the source of DNA and the mechanism of HGT was not determined^35^. Natural transformation may be an important mechanism for the acquisition of antibiotic resistance and virulence genes in this ESKAPE pathogen and a significant contributor to the rapid increase in the number of multidrug resistant *P. aeruginosa* strains that are an emerging problem worldwide.

## Abbreviations

eDNA: extracellular DNA;
T4P: type IV pili;

## Acknowledgements

We thank Jacob Bertrand, Elizabeth Tran, Lisa MacAskill, Dervilla McGowan, Heather Smith and Kate Rainzcuk for technical assistance and helpful discussions. L.M.N was supported by an Imperial College Research Fellowship. D.L. was supported by the Swiss National Science Foundation (SNSF, grant n. P2GEP3_161769). C.B.W. was supported by a National Health and Medical Research Council of Australia (NHMRC) Career Development Award and a Senior Research Fellowship (571905).

## Author information

### Contributions

L.M.N, L.T, M.K, S.R.O, D.L, J.J.L. and C.B.W performed experiments. L.M.N, L.T and C.B.W analyzed data. C.B.W. provided project administration and funding. L.M.N. and C.B.W. wrote the manuscript.

### Ethics declarations

The authors declare no conflict of interest.

## Supplementary Figures

**Supplementary Figure 1.**
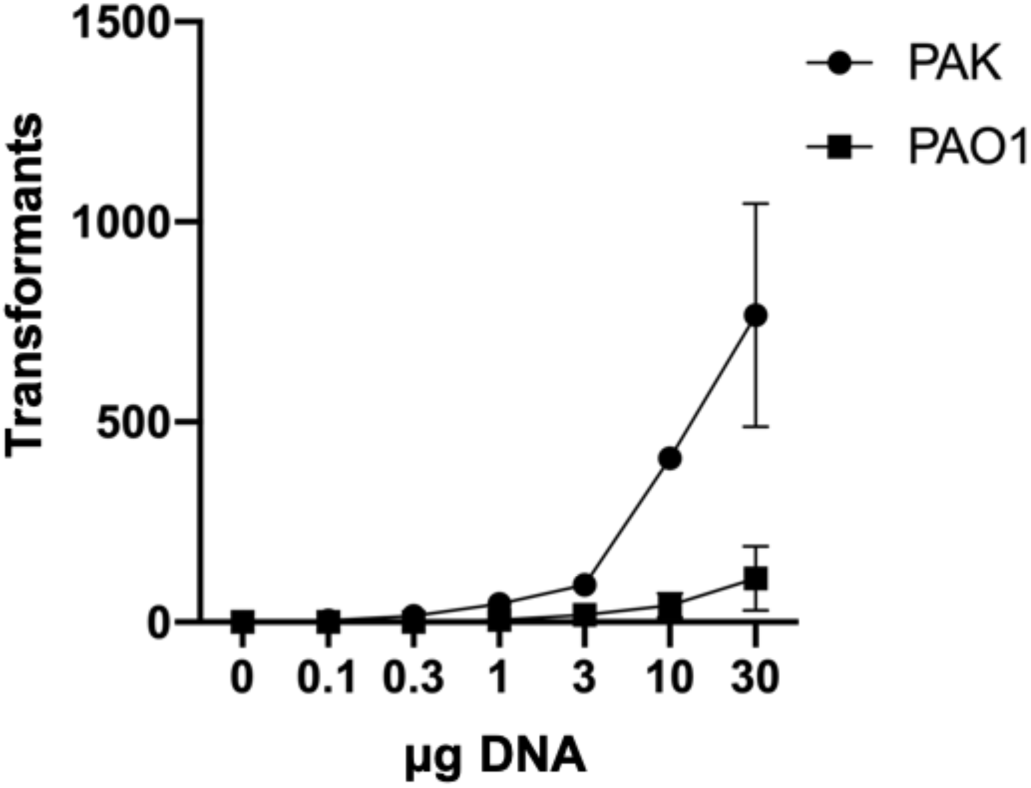
Transformant numbers increase with plasmid DNA concentration in static broth cultures. Static broth cultures of PAK or PAO1 with 10 μg pUCPSK added were incubated and the numbers of carbenicillin resistant transformants determined after 24 hr. The mean of each set of technical triplicates was calculated to give an n≥3 which is presented as mean ± S.E.M.

**Supplementary Figure 2.**
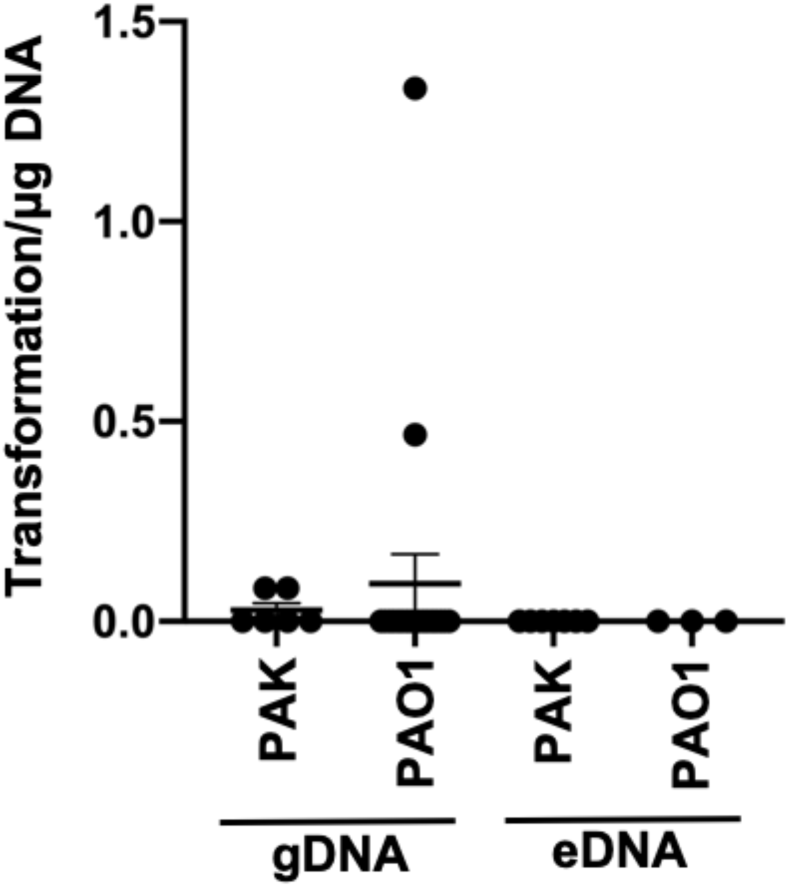
Natural transformation of gDNA and eDNA is inefficient in static broth cultures of *P. aeruginosa*. Static broth cultures of PAK or PAO1 with 15 μg gDNA or eDNA were incubated and the numbers of gentamicin resistant transformants determined after 24 hr. The mean of each set of technical triplicates was calculated to give an n≥3 which is presented as mean ± S.E.M.

